# Streamlined low-input transcriptomics through EASY-RNAseq

**DOI:** 10.1101/564468

**Authors:** Yiwen Zhou, Hao Xu, Haiyang Wu, Haili Yu, Peng Zhou, Xin Qiu, Zihan Zheng, Qin Chen, Fa Xu, Gang Li, Jianzhi Zhou, Gang Cheng, Wei He, Liyun Zou, Ying Wan

**Affiliations:** Biomedical Analysis Center, Army Medical University, Chongqing, China; R&D Dept., TCRCure Ltd., Chongqing, China; Department of Cardiology, PIDU District People’s Hospital, Chengdu, China; Biowavelet Ltd., Chongqing, China; Department of Gynecology and Obstetrics, First Affiliated Hospital of Army Medical University, Chongqing, China; Department of Immunology, Army Medical University, Chongqing, China

**Author notes:** To whom correspondence should be addressed: Ying Wan Biomedical Analysis Center, Army Medical University, Chongqing, China. denotes co-first authors (Yiwen Zhou, Hao Xu, Haiyang Wu).

**Keywords:** T-cells, embryo, mRNA, transcriptomics, low-input, biotechnology

## Abstract

High-throughput sequencing for transcriptome profiling is an increasingly accessible and important tool for biological research. However, accurate profiling of small cell populations remains challenging due to issues with gene detection sensitivity and experimental complexity. Here we describe a streamlined RNAseq protocol (EASY RNAseq) for sensitive transcriptome assessment starting from low amount of input materials. EASY RNAseq is technically robust enough for sequencing small pools of homogenous and heterogeneous cells, recovering higher numbers of genes and with a more even expression distribution pattern than other commonly used methods. Application of EASY RNAseq to single human embryos at the 8-cell stage was able to achieve detection of 70% protein-coding genes. This workflow may thus serve as a useful tool for sensitive interrogation of rare cell populations.

## Introduction

RNA sequencing (RNAseq) has become a potent method for transcriptome profiling, with applications that include monitoring gene expression profiles, discovering and assembling novel transcripts, and investigating alternative splicing. As researchers seek to obtain increasing amounts of information from decreasing amounts of input material, there is an increasing demand for more sensitive and resource-effective sequencing methods. Indeed, the newest wave of techniques has made it possible to generate substantial transcriptomic insights into single nuclei, allowing for a fuller appreciation of the true scope of cellular heterogeneity. These achievements have required the development of novel methods to expand the detectable nucleic acid signal, while balancing it with the loss of fidelity to the original sequences that can occur during the amplification process.

One common approach to overcome the challenge of starting from low amounts of input materials has been to perform whole transcriptome amplification (WTA) reactions to reach the DNA threshold necessary for sequencing library construction. Two major methods for conducting the WTA reaction are in common use. The earliest WTA strategy following the method developed Kurimoto et al. performs a PCR reaction after the polyA tailing in a reverse transcription reaction(1–3). Generally, this strategy would obtain full-length and truncated cDNA. An alternative approach has been to apply a template-switching reaction to ensure production of full-length cDNA(4, 5). However, both WTA strategies are reliant on a PCR reaction, during which unavoidable biases may be introduced to the small amount of input material(6). To reduce these biases, in vitro transcription (IVT) methods have also been developed for use in single-cell RNA-Seq methods such as CEL-Seq and MARS-Seq.(7, 8) Unfortunately, traditional IVT requires intensive laboratory work for the amplification, and is not very easily expandable without automation. These workflows are also time-intensive, leading to increased risk of potential degradation and/or contamination. As such, there remains a need for newer technologies to efficiently assess the transcriptomes of small cell populations.

In this manuscript, we describe a rapid and efficient method to obtain sufficient DNA for sequencing without relying on PCR amplification. Our method is coupled to an accelerated method of RNA isolation that further improves assay sensitivity and processing time. Through comparison of EASY RNAseq performed on input cell counts of varying orders of magnitude, we observed that the procedure was highly robust, and could detect significantly more protein-coding genes than in datasets generated by conventional methods, while efficiently recovering transcripts associated with smaller subpopulations of cells. When applied to two replicate populations of 8-cell stage oocytes, EASY RNAseq could detect >15,000 genes in both samples with high inter-sample correlation. As such, EASY RNAseq is well suited for whole transcriptome profiling of rare cell populations.

## Results

### EASY RNAseq is highly reproducible at low sample input levels

Starting with total RNA, the protocol of EASY RNAseq is divided into five steps (a flowchart of this protocol is provided as Supplementary Figure 1). An initial reverse-transcription reaction is performed using a M-MLV reverse transcriptase with an anchored polyT primer. Next, second-strand synthesis is performed via the combined activity of RNase H and DNA polymerase I, rather than through the conventional PCR-based method. The step performed on small amounts of cDNA allows for EASY RNAseq to be less prone to PCR bias. The resulting samples are then tagmented with Tn5 for adapter ligation and fragmentation, and enriched via PCR prior to sequencing. As the entire EASY RNAseq workflow only requires one PCR reaction, the sequencing libraries for a sample can be constructed within 5 hours.

As a preliminary evaluation of our workflow, we assessed the robustness of EASY RNAseq across a range of starting material levels. Four samples of 5.0 ng, 1.0 ng, 0.5 ng, and 0.1 ng RNA derived from mixed murine splenocytes were generated through serial dilution to generate twelve libraries for sequencing. These libraries yielded a total of 308 million 150-nt paired-end reads, with an average of 28 million reads per library. All sequencing data were analyzed by following a widely used analysis workflow recommended by Sahraeian et al(9) (Table S2). Over 12,000 genes could be found in the 0.1ng sample, representing greater than 60% of the known murine transcriptome (Fig 1A). Despite the differences in the amount of starting materials, libraries constructed by EASY RNAseq protocol were highly similar, with pairwise Pearson’s correlation coefficients exceeding 0.97 after averaging the expression of the three replicates (Fig 1B). Genes with low expression exhibited greater variance, suggesting the possibility that the aliquoting may have led to loss of some rare transcripts (Fig 1C). Notably, nearly all detected genes had a CV less than 1 within replicates of the 5ng sample, with only a slight increase of variance observed in those of the 1ng and 0.5ng samples (47 and 78 genes fluctuating, respectively). From visualization of the alignment of the reads against their gene location, we could see that EASY RNAseq data could generally span across full-length transcripts, with a slight coverage bias favoring the 3’ end of genes (Fig 1D). This 3’ bias is likely caused by incomplete reverse transcription when using polyT primers, and was not unanticipated (10). Encouragingly, less than 1% reads were mapped to rRNA in all results, suggesting that the RNA samples isolated with our protocol were of high quality and efficiently purified. At the same time, a large proportion of data mapped to non-coding region was noticed, including both intronic and intergenic regions. These reads suggest that there may be complex post-transcriptional regulation mechanisms occurring in the cells that has been previously underappreciated, but which EASY RNAseq may be able to successfully detect(11–14).

**Figure 1.**
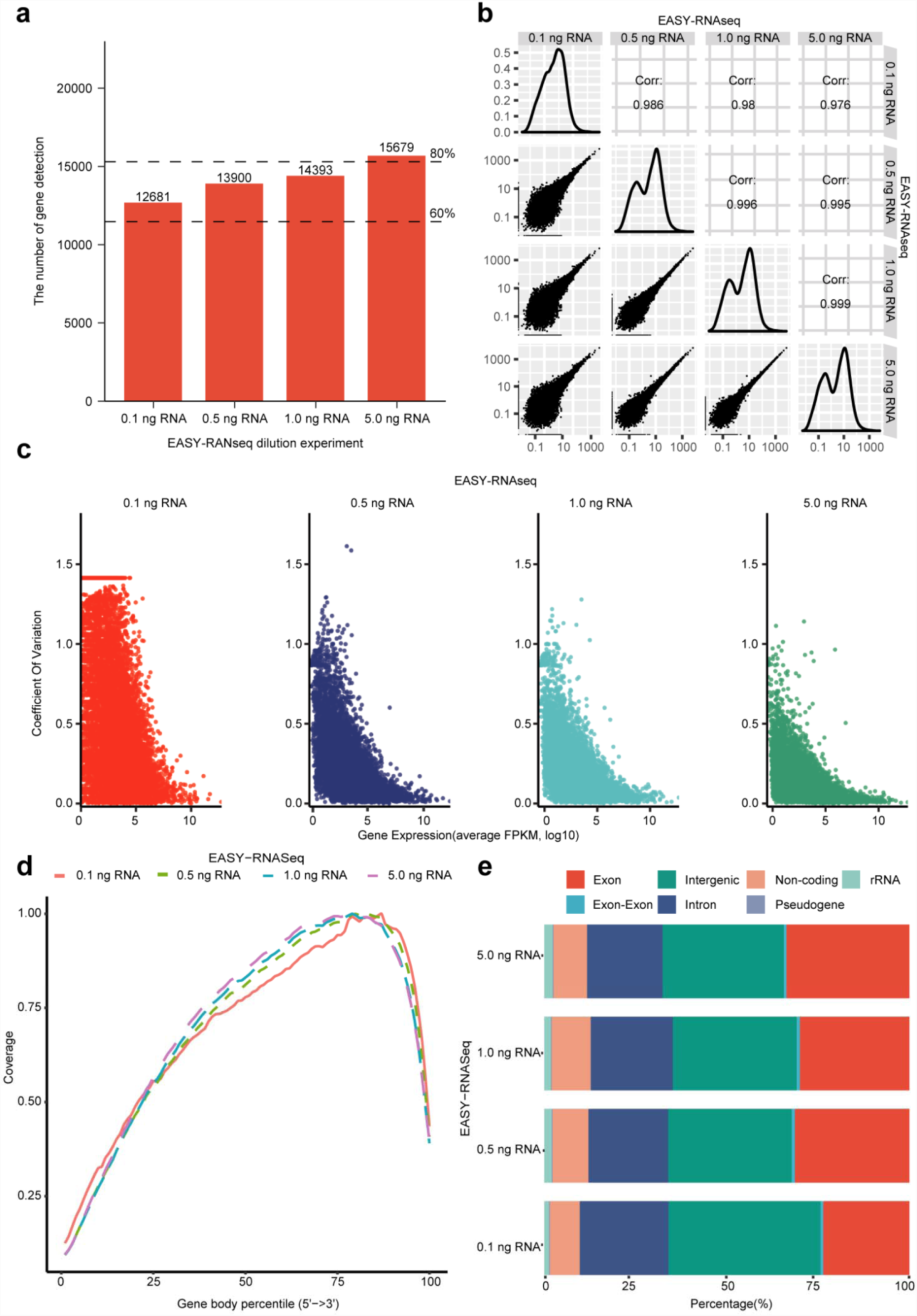
EASY RNAseq is stable for diluted inputs. Serial dilution of from a single RNA sample derived from whole splenocytes could be sequenced consistently using our workflow, with over 12,000 unique genes detected from 0.1 ng of RNA, and increasing based on RNA input, reflecting the increased detection of rare transcripts (A). High sample correlation (B) and low CoV (C) demonstrate that the technique is robust, with appreciable tailing off of COV among highly expressed genes (log10 FPKM >5). Approximately 23% gene body bias was observed that is independent of input quantity (D), and the samples had similar mapping percentages to gene features (E), suggesting that these results are characteristic.

### EASY RNAseq can recover weakly expressed transcripts from small inputs

While the sequencing of aliquoted DNA samples suggested that EASY RNAseq is robust enough to work with small input material, most biological questions instead involve starting from small populations of cells. As such, we next generated individual samples of sorted T cells, and compared these results back to our preliminary whole-splenocyte mixtures. Consistent with our expectations, the vast majority of genes identified in the sorted T cells had been recovered in the splenocyte samples (Fig 2A). Visualization of the variably expressed genes demonstrated that there was a clear skew towards higher expression in the whole splenocyte samples, including of myeloid and B cell genes such as Cd33 and Jchain (Fig 2B), suggesting that EASY RNAseq could effectively recover transcripts from subpopulations within a sample. This observation was further supported by gene set enrichment analysis (Fig 2C) and pathway analyses (Fig 2D) in which B cell associated pathways were highly enriched in the splenocyte samples. At the same time, we could clearly observe transcripts associated with T cell status and of signature transcription factors associated with smaller T helper subpopulations (Fig 2E) even in the smallest 100 cell samples. Deconvolution analysis of the samples could also identify a clear enrichment for non-T cell types in the splenocyte samples (Fig 2F). These results confirm that EASY RNAseq is sufficiently robust to recover meaningful information about subpopulations of cells.

**Figure 2.**
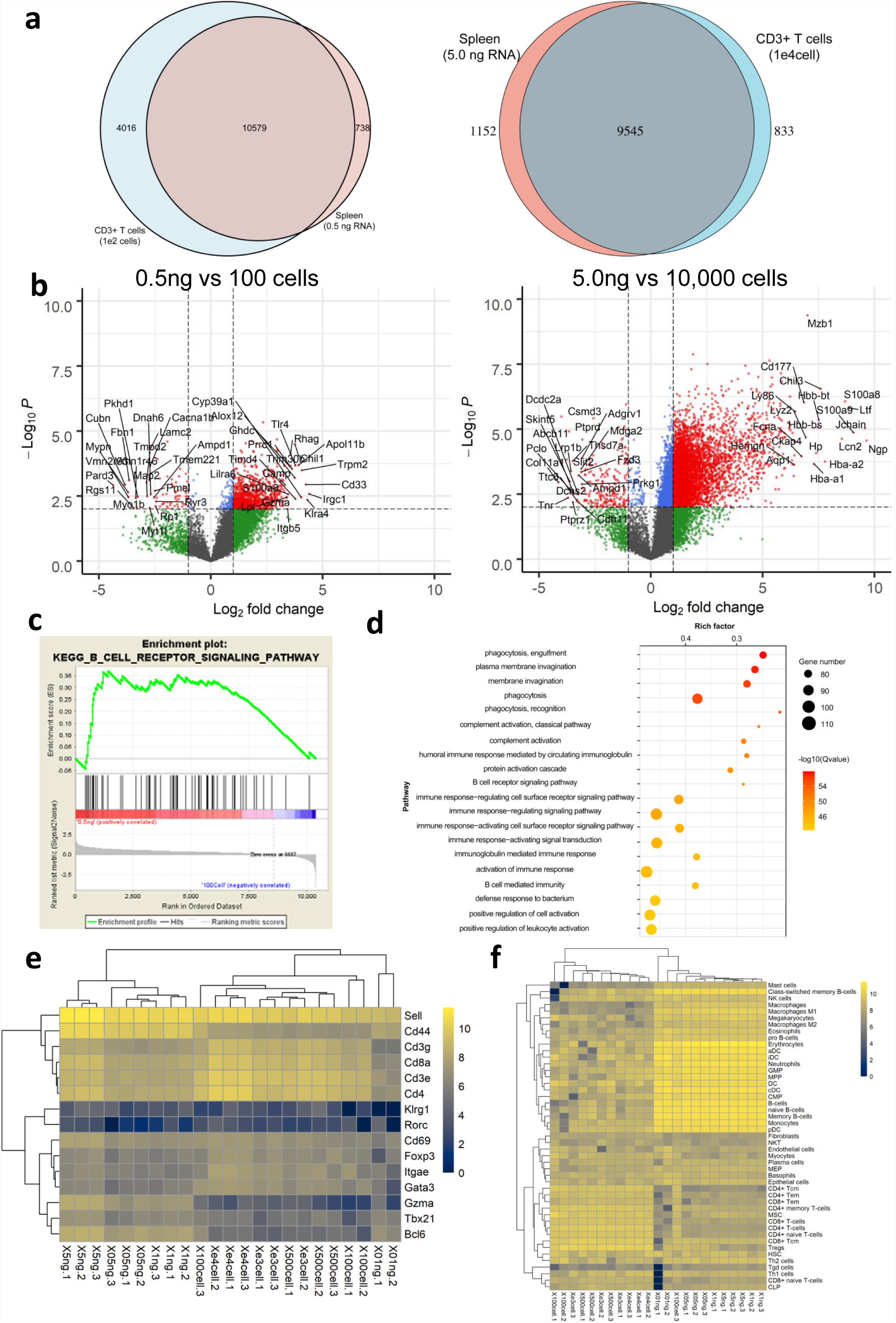
Recovery of subpopulation transcripts in heterogeneous mixtures. Comparison of the splenocyte and purified T cell samples demonstrates that the large majority of the genes recovered between the two are the same, regardless of relative input amount (A). (B) Volcano plot visualizations show significant skew in favor of higher expression in the splenocyte samples, with 1,500 of the genes showing a difference greater than 2 between the 5ng and 1E4 samples. (C,D) Representative GSEA plot and pathway analysis uncovers a number of pathways associated with B cells and macrophages. (E) Heatmap of signature T helper subset transcription factors (Foxp3, Rorc, Tbx21, Gata3, and Bcl6), T cell status markers (Cd69, Cd103, Cd62L, Cd44, Klrg1) also finds an increased expression of Cd3e, Cd3g, Cd4, and Cd8 in the purified T cell samples consistent with our expectations. (F) Deconvolution analysis is able to identify the presence of dendritic cells, granulocytes, and progenitors that are minor populations within the spleen.

In order to better assess the detection capability of EASY RNAseq, we further matched our sequencing results with public T cell sequencing data generated via other methods. Comparison of gene counts revealed that EASY RNAseq was able to detect over 14,000 unique genes from just 100 input cells, exceeding the amount recoverable from the use of other protocols (Fig 3A), while over 17,000 genes could be recovered from 10,000 cells. This effect was even more striking when the results were downsampled to an equivalent number of input reads. Indeed, the saturation curve of the EASY-RNAseq sample very quickly surpasses the others in gene count, but appears to approach saturation more slowly (Fig 3B). Since the quantified expression of any individual gene in RNAseq is generated through competition to be sequenced, one of the main draws/difficulties of RNAseq is its ability to evaluate relative expression of different transcripts within a single sample. Violin plots of the distribution of gene expression illustrated that EASY RNAseq profiled gene expression with a more centered density, suggested that EASY RNAseq could recover both more weakly expressed transcripts and find them at higher absolute FPKM (Fig 3C). At the same time, the genes that were detected had high expression correlation when compared with the other datasets, except for the SMARTseq set that sequenced regulatory T cells (Fig 3D), indicating that our sequencing approach was still reliable. EASY RNAseq was able to successfully quantified 4626 more genes at any level, of which 1302 could be found at FPKM greater than 1 (Table S3, Fig 3E). To further verify those genes were true positives, the frequency distribution of relative expression was split into two categories: all detected genes and the uniquely detected genes. As expected, the distribution of unique genes expression was centered at low expression values, with the few genes missed in EASY RNAseq data also being ones with relatively low expression levels (and potentially attributable to random sampling bias) (Figure 3F).

**Figure 3.**
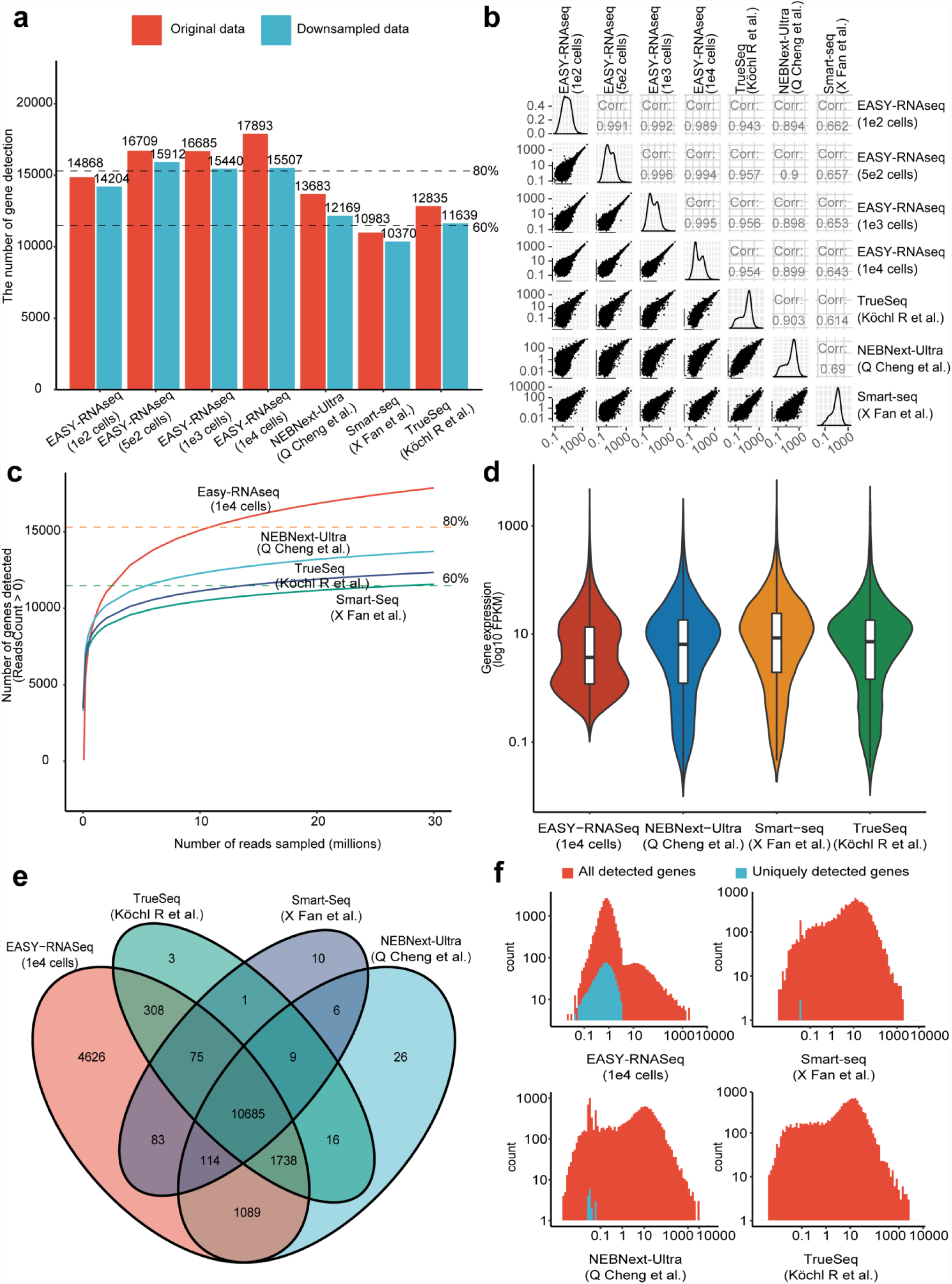
Comparison of EASY RNAseq with other sequencing methodologies. (A,B) Downsampling of each sequencing run to 1E7 reads and full comparisons demonstrate that EASY RNAseq is able to more evenly recover unique genes than other sequencing workflows, but still has very high correlation with the data generated by other methods in terms of expression levels. (C,D) Saturation analysis and violin plots of the expression distribution of each method reveals a notable bulge in genes with low expression that is not present in the other datasets, and is also shifted upwards with a higher minimum. (E,F) Venn diagram of the unique genes found by each method, and the expression levels of the uniquely detected genes within each method shows that most of the unique genes still have expression above 0.1 FPKM in EASY RNAseq.

### EASY RNAseq profiling of embryos

Having verified that EASY RNAseq is sufficiently robust for transcriptome analysis of low input mouse samples, we next sought to apply the approach to even smaller counts of human cells as another likely use scenario. RNA from two pairs of twelve cells harvested from rejected IVF embryos were processed using the EASY RNAseq workflow, leading to recovery of 21.48 million reads. 82.86 % and 90.35 % of the reads mapped to reference genome, with 55% of them being exonic in the former and 43% exonic in the latter (Table S4). Interestingly, the difference of exonic mapping percentage did not seem to affect the overall gene detection sensitivity, as those exonic data detected 15,191 and 15,048 genes, respectively. Among them, 10,893 genes were quantified with at least 5 reads in both replicates, accounting for 83% of all detected genes (Fig 4A). The ranked correlation of the two replicates was 0.88, and most of the genes that failed to match between the two were of extremely low expression, suggesting that the protocol was able to capture the majority of biologically significant information (Fig 4B). While functional analyses of the top 1000 expressed genes unsurprisingly unveiled high enrichment for genes associated with basic cellular machinery (Fig 4C), comparison with a previously published set of genes varying during early embryonic development (35) confirmed that a range of randomly selected high and low expressed genes with our data had the same general expression pattern (Fig 4D). Indeed, integration of our sequencing data with a previously published 8-cell stage dataset (36) revealed that 12,415 genes were found to be shared between our two samples and the previous data (Fig 4E). Correlational analysis demonstrated that our samples had a 0.7 correlation with that set (Fig 4F). Collectively, these results demonstrate that EASY RNAseq is suitable for detailed transcriptomic investigations of rare/small cells populations.

**Figure 4.**
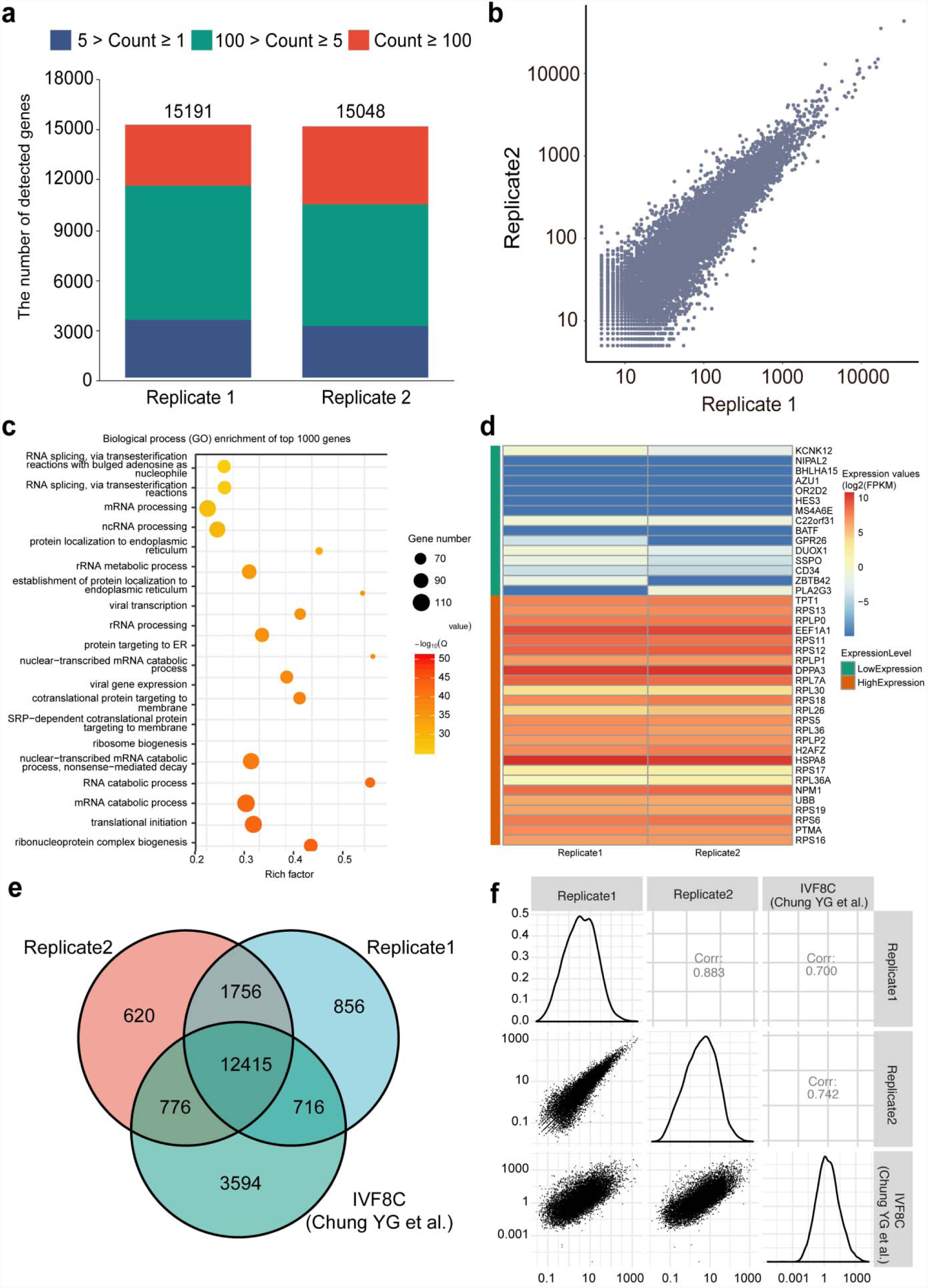
EASY RNAseq can efficiently sequence 8-cell embryo samples. (A,B) The majority of genes in both samples were detected at robust expression levels, matching the distribution expected based on our previous sequencing, while also sharing high correlation. (C,D) Functional analysis and gene list comparison confirms that our sequencing data encompasses expected biological activities and factors previously identified. (E,F) Comparison of our samples with a previously published report on 8-cell stage embryos shows that our approach is able to identify a similar number of common genes, as well as a significant amount of unique factors. Spearman’s ranked correlation of the samples confirms that the genes are expressed in similar patterns in both.

## Discussion

The true breadth of cellular heterogeneity and the significance of small populations of cells is becoming increasingly appreciated as new technologies for assaying larger amounts of parameters per cell have been developed. While traditional flow cytometry and imaging approaches are typically limited to under 10 parameters, new methods such as mass cytometry and multiplexed immunofluorescence have expanded the detectable repertoire to achieve over 40 parameters. This increase in observable variables has sharply expanded due to the advent of single-cell RNAseq, with advanced platforms now capable of identifying up to 8,000 genes per cell in hundreds of cells. Despite these advances however, the need for accurate transcriptome measurements of small mixtures of cells has not gone away. Due to gene dropout and other technical limitations, most approaches for differential analysis of gene expression in single-cell data requires for the *in silico* clustering of the most similar cells, and are critically dependent upon bulk RNA sequencing for validation. Our recovery of over 13,000 shared genes from our application of EASY RNAseq on human embryos as the 8 cell stage represents a much higher detection rate than what is currently achievable by single-cell techniques, and could thus serve as a useful scaffold for validating single cell results. Indeed, it has been suggested that the actual RNA content of embryos at this stage may be less than that recoverable from a single unfertilized oocyte (34). By avoiding the bias induced by the PCR step prior to cDNA synthesis in conventional WTA, EASY RNAseq is able to more accurately recover transcripts while also being time efficient and highly reproducible. The simplification offered by EASY RNAseq may also allow it to be more amenable for automation, as most steps are reduced to simple liquid handling.

Successful transcriptome profiling of rare cell population requires overcoming the hurdle of needing to amplify low input materials while insulating against technical noise. To meet these criterion, EASY RNAseq applies a PCR-free method to replace the widely used whole transcriptome amplification for second strand synthesis. With only a dozens of cells, EASY RNAseq is capable of detecting more than 15,000 of protein-coding genes. Among these detected genes, nearly 80% entries have strong signals supported by more than 5 independent reads. This level of sensitivity can greatly enhance both sensitivity and accuracy of distinguishing subgroups from rare cells, and aid in feature selection of cell subsets for development of novel biomarkers. EASY RNAseq may also allow for easier application of deconvolution techniques for characterizing subpopulations of cells within pooled populations as a result of the advantages offered by the more linear distribution pattern of transcript expression compared to other existing techniques. Furthermore, while transcriptome analysis is best performed on fresh samples, clinical samples are often preserved for extended periods. The ability of EASY RNAseq to successfully sequence starting from low amounts of heterogeneous input suggests that it may also be capable of sequencing RNA from rare clinical specimens. Future explorations and refinements are still necessary to clarify the full potential of this approach.

## Materials and Methods

### Sample preparation

C57BL/6 female mice were obtained from Beijing Huafukang Bioscience and maintained under specific-pathogen-free (SPF) conditions. All animal experiments were conducted according to protocols approved by the Medicine Animal Care Committee of the Third Military Medical University. The spleen were isolated with dissecting forceps gently from the 8-week-old C57/BL6 wild-type female mouse after sacrifice with carbon dioxide, and grinded into a single cell suspension with syringe piston through the 100 µm strainer. Erythrocytes were cleared by resuspending the cells in ACK lysis buffer (Tiangen) per manufacturer’s protocol. The splenocytes (1e6) were stained with Anti-CD3e-Percp-cy5.5 (BD Pharmingen, Cat. 551163) for 25 min of staining on ice, and then with DAPI for dead exclusion. Samples of CD3 positive T cells (1e2/5e2/1e3/1e4) were sorted by flow cytometry with the BD Jazz. All human oocyte samples for EASY RNAseq profiling were obtained after in vitro fertilization, and had been discarded following observation of tripronuclear (3PN) cells in the zygotes at Day 1. Use of these samples was permitted by the Ethics Committee of the First Affiliated Hospital, Third Military Medical University under Approval No. 201554 to W.H.

### RNA acquisition

All cell samples were stored in TRIzol reagent (Invitrogen, cat. no. 15596026) at −80 °C prior to RNA isolation. Bulk quantities murine RNA was isolated by using classic liquid phase separation methods with trichloromethane and isoproply alcohol(15, 16), and then diluted to 10 ng/µl for each sample. The normalized sample was further divided into different starting material amounts (0.1/0.5/1.0/5.0 ng) by using gradient dilution. RNA was isolated from different number of murine splenic CD3 positive T cell samples and human oocyte samples by using the Direct-zol™ RNA MiniPrep Kit (ZYMO, cat. R2050). The extracted RNA samples were dissolved into 30 µl of nuclease-free water (Qiagen, cat. 129115). RNA concentration and the absorbance ratio at 260/280 nm and 260/230 nm of each sample was measured by NanoDrop™One (Thermo, cat. ECS000493), with a small aliquot of each used to assess 28S/18S by gel electrophoresis for further quality control.

### EASY-RNAseq Library preparation

The qualified RNA samples were added with 1 µl of SuperScript III reverse transcriptase (200 units/µl) [Invitrogen, cat. 18080044], 4 µl of Superscript III first strand synthesis buffer (5x) [Invitrogen, cat. 18080044], 1 µl of dNTP mixture (10 mM), 1 µl of Oligo 30 (dT) primer (10uM) [polyT primer], 1 µl of RNase inhibitor (40 units/µl) [Thermo, cat. k1622] and 1 µl of DTT (0.1M) to construct 20 µl reaction system for RNA reverse transcription(17) by heating the mixtures for 50 °C for 90 min, 70 °C for 15min, and a final hold at 4 °C. For second strand cDNA synthesis, the 20 µl products of reverse transcription were directly mixed into reaction buffer (NEB, cat. E6111S/L) on ice to obtain dsDNA samples with the help of DNA polymerase I and T4 DNA ligase, following the manufacturer’s recommendations (18, 19). dsDNA products obtained from the synthesis were purified with 144 µl (1.8 ×) of Agencourt AMPure XP Beads (Beckman, cat. A63880-A63882) through magnetic separation (Invitrogen, cat. no. 123.21D)(19). The purified dsDNA samples were then dissolved into nuclease-free water and diluted to 1 ng/µl following Qubit 2.0 measurement (Invitrogen, cat. Q32866). 1 ng of dsDNA, and 4 µl of 5 × TTBL and 5 µl of TTE Mix V1 were used to in the Tn5 tagmentation reaction system. Reaction was performed for 10 mintues at 55°C and quenched with 5 µl of pre-mixed 5 × TS to avoid excessive DNA fragmentation. Samples were then barcoded following the protocol of TruePrep DNA Library Prep Kit for Illumina with P5/P7 adapter primers (Vazyme, cat. TD503), and amplicon purified via the VAHTS™ DNA clean beads kit (Vazyme, cat. N411-02).

### Analysis of RNA-Seq data

All EASY-RNAseq libraries were sequenced through Illumina HiSeq platform with 150bp pair-end model. Three published datasets were used for method comparison (GEO Accession: GSE63961, GSE121482, GSE111066). All sequencing reads were passed through adapter filtration by using Trimmomatic(20). The clean data were aligned to respective genome reference (GRCm38 and GCRh38 respectively)(21). Mapped reads uniquely assigned to one genomic location and one gene were counted as gene expression, which was carried out by FeatureCounts(22). Normalization of gene expression was performed by transferring the read counts to FPKM values. And both of the read counts and FPKM values were used in visualization. The coverage of gene body and percentage of data features were calculated by RseQC(23). Those downsampled data were directly made from corresponding raw data and analyzed by same protocol.

The reproducibility was assessed within samples and replicates respectively. Pearson’s coefficient was used to measure the similarity between samples, which was calculated with average FPKM values. Similarly, the Coefficient of Variance(CV) was applied to represent the technical robustness within replicates. The saturation curves were created by the regression of downsampled datasets. Each curve was based on 25 different volumes of downsampled data, ranging from 0.1 million to the corresponding original data size. The saturation plot was cutoff that only showed the maximum sequencing depth of 30 million. The Venn plot was draw with those quantified (reads count > 0) protein-coding genes of each original dataset. The uniquely quantified genes of each dataset were selected, together with all quantified protein-coding genes, were visualized as expression frequency spectrums. Gene set enrichment analysis was performed using the GSEA app (Broad Institute) on the KEGG gene lists. Deconvolution analysis was performed using XCell in R according to default parameters (33).

### Transcriptome profiling of embryonic cells

The expression values of two embryonic samples were obtained by the same RNA-Seq analysis workflow mentioned above. The expression values were defined into three categories for visualization. Among those quantified genes, genes with reads count greater than or equal to 5 were treated as confidently detected. To decrease the false positives, only those confidently detected genes were used in following analysis. The Gene Set Enrichment Analysis (GSEA) was performed with the intersection of expression results through clusterProfiler packages(24, 25).

### Visualization

All figures were produced by R(26) and ggplot2(27). The color board was using ggsci(28). Among them, those plots of Pearson’s coefficient in Fig2.b and Fig4.b were created by GGally(29). And the Venn plot was produced through VennDiagram(30). Volcano plots were visualized through EnhancedVolcanoplot (31) and heatmaps through pheatmap (32).

## Acknowledgments

This work was supported by grants from the National Key Research and Development Program of China (2017YFA0700404) for Y.W., and the National Natural Science Foundation of China (91642119) for Y.W. We would like to thank the other members of the Wan lab and Dr. Lin He for insightful discussions during the drafting of the manuscript.

## Potential Conflicts of Interest

Y.W., L.Z., and J. Z. are affiliated with Biowavelet Ltd, which is developing automated methods for performing biological experiments.

**Supplementary Figure 1.**
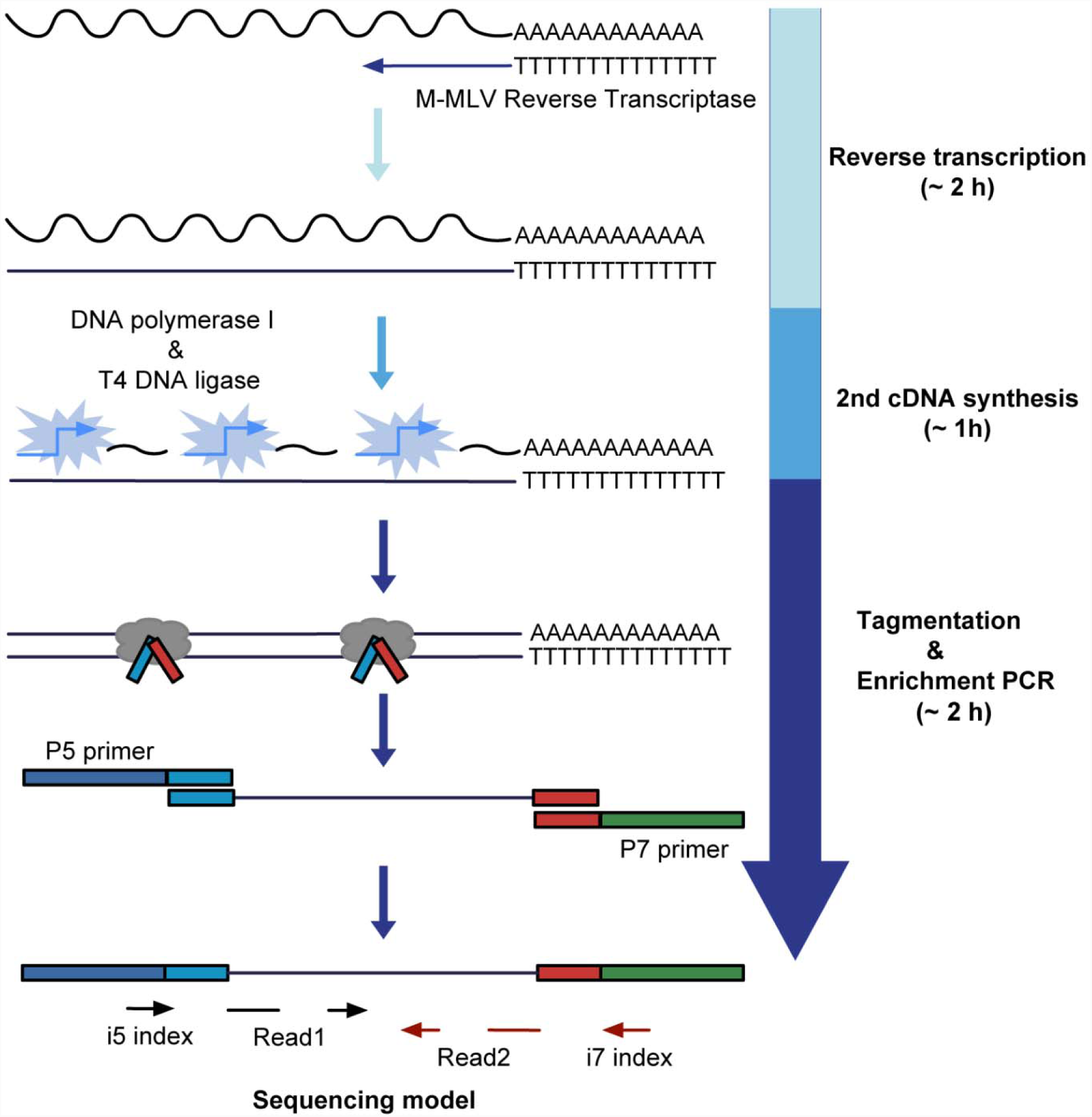
EASY RNAseq library construction workflow.

