## Supplementary figures and images for "Streamlined low-input transcriptomics through EASY-RNAseq"

### Supplementary Figure 1

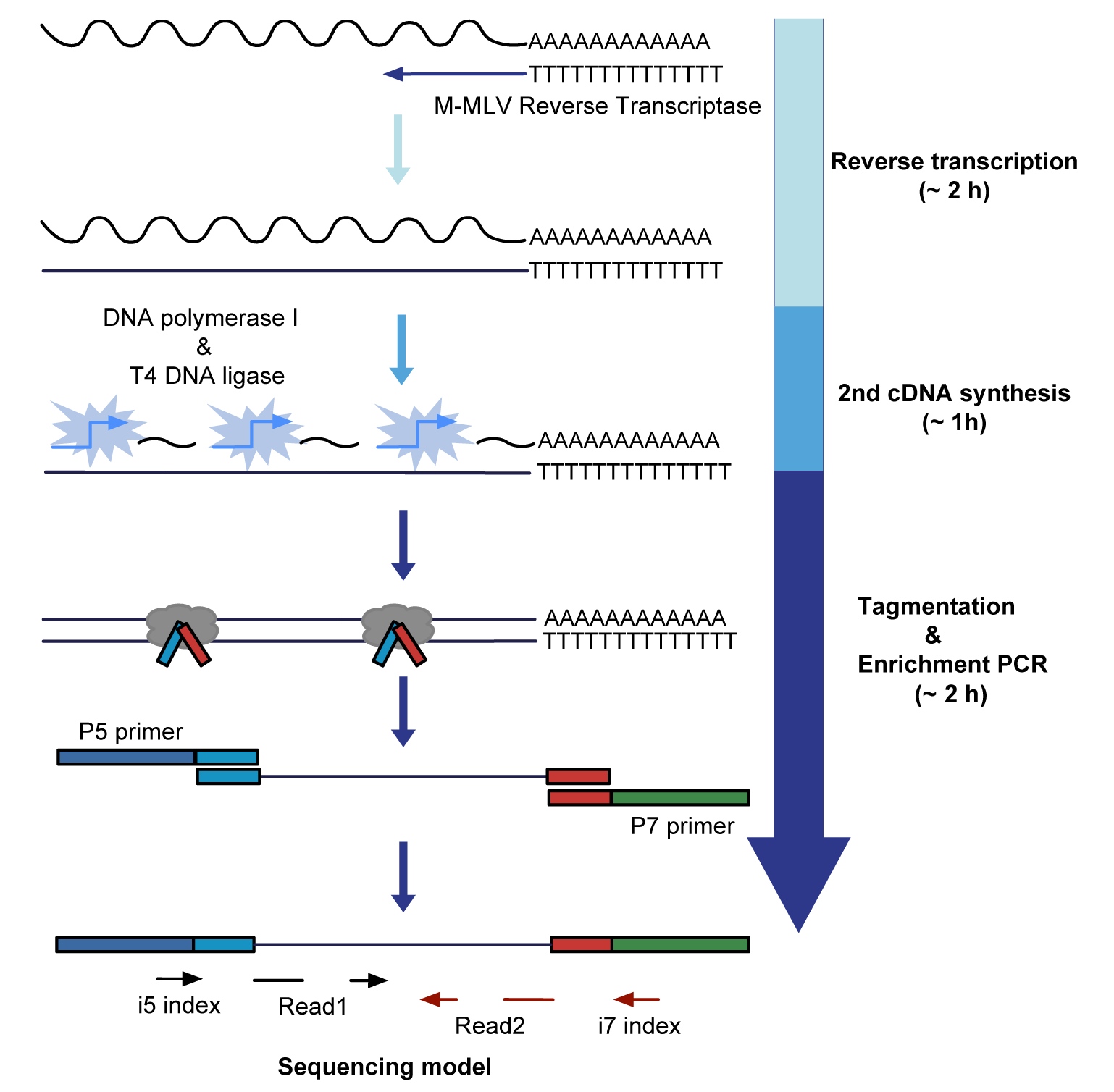
